# Non-coding 886 (*nc886*/*vtRNA2-1*), the epigenetic odd duck – implications for future studies

**DOI:** 10.1101/2023.09.29.560139

**Authors:** Emma Raitoharju, Sonja Rajić, Saara Marttila

## Abstract

Non-coding 886 (*nc886*, *VTRNA2-1*) is the only human polymorphically imprinted gene, in which the methylation status is not determined by genetics. Existing literature regarding the establishment, stability, and consequences of the methylation pattern, as well as the nature and function of the nc886 RNAs transcribed from the locus, are contradictory. For example, the methylation status of the locus has been reported to be stable through life and across somatic tissues, but also susceptible to environmental effects. The nature of the produced nc886 RNAs has been redefined multiple times and are still under debate and in carcinogenesis, these RNAs have been reported to have conflicting roles. In addition, due to the bimodal methylation pattern of the *nc886* locus, traditional genome-wide methylation analyses can lead to false-positive results, especially in smaller datasets.

Here, we aim to summarise the existing literature regarding *nc886*, discuss how the characteristics of *nc886* give rise to contradictory results, and reinterpret, reanalyse and, where possible, replicate the results presented in the current literature. We also introduce novel findings on how the *nc886* methylation pattern distribution is associated with the geographical origins of the population and describe the methylation changes in a large variety of human tumours. Through the example of this one peculiar genetic locus and RNA, we aim to highlight issues in the analysis of DNA methylation and non-coding RNAs in general and offer our suggestions for what should be taken into consideration in future analyses.

## Background

Non-coding 886 (*nc886*, HGNC symbol *vtRNA2-1*, previously referred to also as *pre-miR-886*, *CBL3* and *hvg-5*) is the only known polymorphically imprinted gene in humans, the variation of which is not caused by genetic factors^1–5^. In population cohorts consisting mainly of Caucasians, 75% of individuals have been reported to present a maternally imprinted region in this locus (chr5:136078784-136080957, GRCh38^2^), while approximately 25% of individuals present two non-methylated alleles^1,6–8^ (Figure 1A). A non-coding RNA is transcribed from the locus, but the nature of the RNA is still largely under debate, i.e. whether the full-length RNA forms a hairpin structure and bounds directly to proteins or is cleaved into miRNA-like short RNAs^9,10^. The expression of this RNA is regulated by the methylation pattern in the gene locus^1,11,12^, but also by the genetic variation down-stream of the gene^1^.

**Figure 1.**
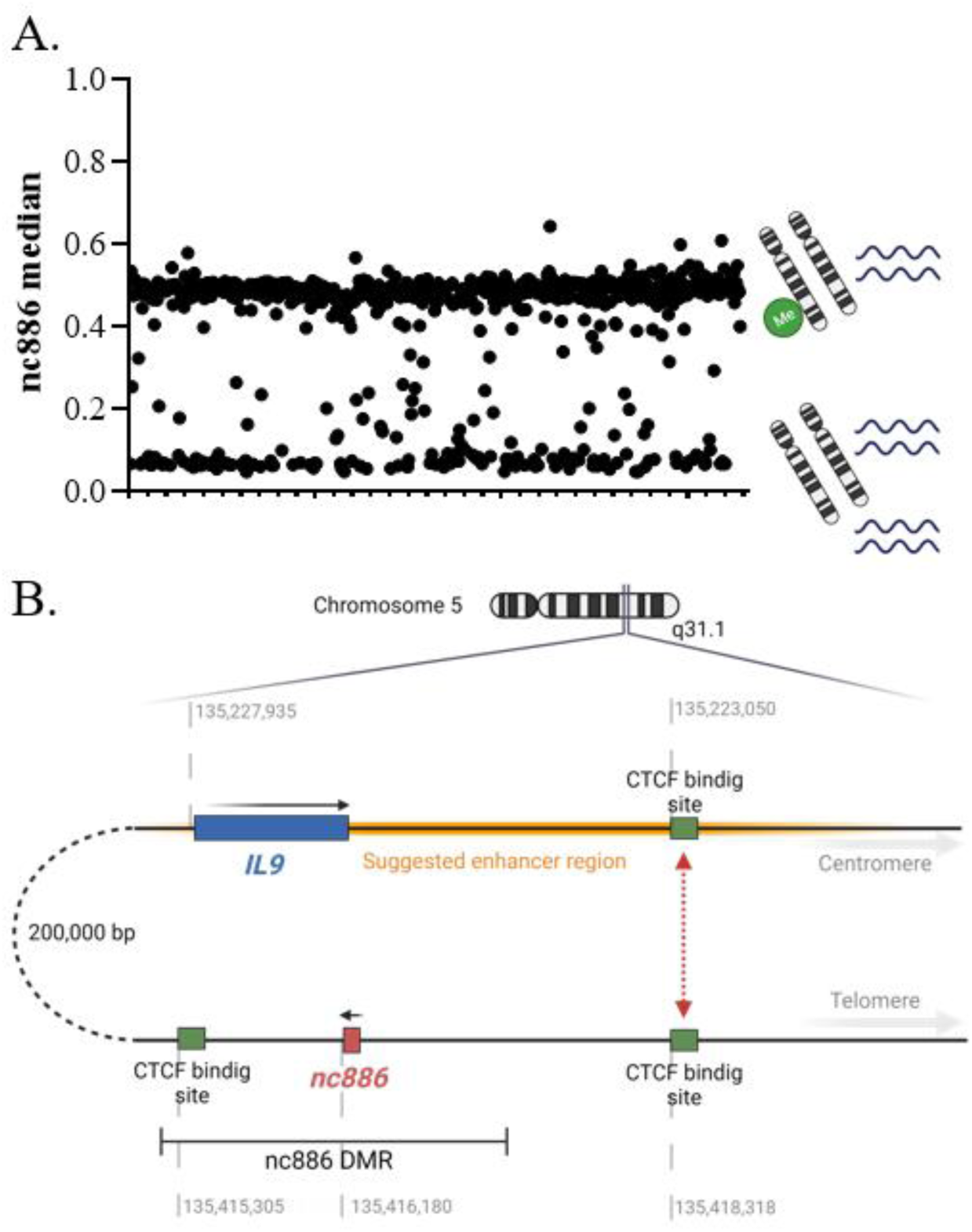
A) Distribution of median methylation levels of cg07158503, cg11608150, cg06478886, cg04481923, cg18678645, cg06536614, cg25340688, cg26896946, cg00124993, cg08745965 and cg18797653 locating in the DMR overlapping *nc886* from Caucasian and Hispanic individuals (GSE40279). Two clusters can be observed: individuals with 50% methylation level (∼75% of the population) and individuals with a methylation level close to 0% (∼25% of the population). In the former, the maternal allele is methylated^6–8,41,42^, while the paternal allele is unmethylated and permissive for transcription, whereas in the latter, both alleles are unmethylated and permissive for transcription. B) Schematic presentation of the *nc886* gene, *nc886* DMR and the CTCF-binding sites flanking the DMR. The telomeric CTCF-binding site has been suggested to interact with another binding site near the *IL9* gene, bringing a suggested enhancer region close to the *nc886* gene^1,27^. Made with BioRender.com.

There are a number of studies showing how preconceptional or prenatal conditions associate with the methylation pattern of this gene^1,7,8,13,14^ and both the methylation pattern and RNAs expressed from this locus have been linked to health traits^1,7^ and morbidities^15,16,16^, making it a candidate for a molecule mediating the Developmental Origins of Health and Disease (DOHaD) hypothesis^17^. However, due to the bimodal DNA methylation pattern of the *nc886* locus and the considerable and stable physiological variation in the nc866 RNA expression, conservative analysis methods may produce false positive or inconsistent results. Here, we discuss why and how the characteristics of *nc886* give rise to the contradictory results presented in current literature, as well as provide new, reanalysed and replicated data. We pinpoint the unanswered questions regarding *nc886* and emphasise the requirements for future studies regarding this genetic locus and the RNA(s) produced from it.

### Discovery of nc886 RNA(s)

Non-coding 886 was first discovered as miR-886-3p and 886-5p by human short RNA sequencing^18^. It was soon identified also in rhesus macaques^19^ and included in miRbase version 10. For an RNA to be considered as a miRNA with high confidence, it should produce a mature product approximately 22nt long, have a hairpin structured precursor, be phylogenetically conserved, and the decrease in Dicer enzyme function should lead to increased amount of precursor molecule levels^20^. These requirements were met by the 108nt long nc886 primary transcript which is cleaved into two short RNAs. In 2009, nc886 RNA was, however, shown to have significant sequence similarities with vault-RNAs^9,21^ and co-sediment with intact vault particles^22^, thus leading to the removal of miR-886 from miRbase (version 16). The RNA was suggested to be a novel vault RNA (vtRNA)^23^ and renamed as *vtRNA2-1*, which remains the HGNC accepted official name of the gene. Like the three previously known vault RNAs (vtRNA1-1, 1-2 and 1-3), nc886 is coded in the 5q31, has a suitable length for a vault RNA, and has been suggested to form similar secondary structures as the other members of the vault family^24^.

Since then, the identity of nc886 transcript both as a miRNA and as a vault RNA has been questioned. In 2011, Lee et al. recharacterised the nc886 RNA and concluded that the stem-loop of nc886 has qualities distinct from classical pre-miRNA and, unlike the majority of pre-miRNAs, its production is independent of Dosha. Dicer was also reported to cleave the stem-loop to two ∼20nt RNAs with poor efficiency and the levels of these short RNAs were shown to be so low that they could not be detected in the majority of the analysed samples^25^. nc886 RNA was also shown to be transcribed by RNA polymerase III^26^, while the majority of human miRNAs are transcribed by RNA polymerase II. On the other hand, it was shown that the sequence similarity to the previously known vtRNAs is mostly limited to regions relevant to transcription by Pol III, the expression pattern of nc886 differs from other vtRNAs, and that vault proteins and nc886 RNA do not to co-localize^25^. The Dicer-dependent, but Dosha-independent production of the small RNAs derived from this region was later verified by Miñones-Moyano et al. and Fort et al. ^9,16^. However, Fort et al. demonstrated that the produced small RNAs associate with Argonaute, satisfying the attributes of miRNA, and further hypothesised that the suggested recent evolutionary origin of the *nc886* gene would be the reason for the poor efficiency of Dicer to produce the mature miRNAs^9,27^. This work was again followed by the work of Lee et al. describing the issues arising from the classification of nc886 as miRNA and further emphasising their previous view of nc886 being distinct from pre-miRNAs^10^.

Regardless of the classification of nc886, the gene has been shown to code for a 108nt long non-coding RNA, transcribed by Pol III^26^, which is then cleaved into two short RNAs (nc886-3p; 23nt, and nc886-5p; 24-25nt) with rather low efficiency^9,25^. The full-length nc886 is highly expressed in at least cancer cell lines (10^5^ copies per cell in HeLa cells), the transcripts are localized in the cytoplasm in an organized way^25,28^ and the half-life of the transcripts have been reported to be short (71.5 min)^12^.

### Function of nc886 RNAs

The function of the nc886 RNAs has been thought to be mediated either by the short RNAs acting as miRNAs^1,9^ or by direct binding of the hairpin structure to protein kinase R (PKR) (nucleotides 46-59)^29–31^ or to 2–5-oligoadenylate synthetase 1 (OAS1)^32^. The direct binding to PKR has been shown to lead to tumour-suppressive consequences^25^ and Lee et al. have suggested a tumour-suppressive model where the binding of nc886 RNA prevents PKR from being activated^33^. On the other hand, the short nc886 RNAs have been repeatedly predicted to regulate pathways linked with cancer development and insulin signaling^1,9^. It has also been suggested that the effects of nc886 RNA would be caused at least partly by its binding to RISC and the subsequent hindering of the processing of other miRNAs^28^. The expression levels of either the full-length nc886 RNA or the short derivates have been associated with infection ^23,34–36^, allergy^37^ asthma^38^ and skin senescence^39,40^. Consequences of nc886 RNA expression have been studied little outside cancer (reviewed later), and many open questions remain about its function in physiological conditions.

### Methylation status of *nc886* locus

The *nc886* gene is located in the only known canonically polymorphically imprinted region, the methylation status of which is not associated with genetics in humans^1–3,5^. In detail, there is an approximately 1,600 bp long differently methylated region (DMR) including the *nc886* coding sequence, which is flanked by two CTCF binding sites, hypothesised to insulate the DMR (Figure 1B)^2,7^. Somatic diploid cells present each genomic site twice (maternal and paternal copies), and as in one DNA strand the methyl group can either be present or lacking, the DNA methylation of a CpG site in one cell can be either 0, 50 or 100%. When analysing samples with a mixture of cells, the DNA methylation of a given CpG site usually becomes a continuous variable, with values ranging from 0-100% (or from 0 to 1 in beta-values), as the sample is comprised of different proportions of cells with the previously mentioned DNA methylation statuses. The DMR overlapping the *nc886* however presents a mostly bimodal methylation pattern, where 75% of individuals have 50% methylation in the region (thus forward referred to as imprinted individuals) and approximately 25% of individuals have methylation levels close to 0 (thus forward referred to as non-methylated individuals), indicating two non-methylated alleles^1,6–8^ (Figure 1B). Studies on family units^41,42^ and gametes^6–8,42^ strongly suggest that it is the maternal allele that is methylated in individuals with a 50% methylation level in this locus. It must be highlighted that the bimodal methylation pattern poses challenges both to data processing^43^ and to analysis, as the majority of genome-wide analyses rely on linear regression methods and thus are based on the assumption that DNA methylation in a given site is a continuous, normally distributed variable.

Due to the parent-of-origin-dependent methylation pattern, *nc886* is considered to be an imprinted gene. Canonically imprinted genes present a parent-of-origin-dependent gene expression pattern, with either maternal or paternal locus silenced via epigenetic mechanisms, including DNA methylation^44^. Typically, imprinted genes present a methylation level of 50% in all somatic cells and tissues^45^. The epigenetic profiles maintaining the imprint are established during gametogenesis when the existing DNA methylation pattern is first erased, and then the parent-of-origin-related DNA methylation pattern is created^46^. This pattern is then retained throughout an individual’s life. In addition, tissue or developmental stage-specific imprinting can be observed, for example, in the placenta^47,48^. The significance of intact genetic imprints is highlighted by the severe disorders caused by imprinting defects^49^. We have previously shown that individuals presenting multilocus imprinting disturbances also present altered methylation pattern at the *nc886* locus^6^, indicating that there are similarities in imprint establishment and/or maintenance of the *nc886* imprint and the more typical non-polymorphically imprinted genes.

Similarly to canonically imprinted genes, individuals present the same methylation pattern of *nc886* DMR in the majority of their somatic tissues, regardless of the germ layer from which the tissue originates^6,8,13^. We have previously reported that among the 30 studied somatic tissues, only skeletal muscle and cerebellum make an exception. In skeletal muscle, all individuals present an imprinted *nc886* profile with a 50% methylation level and, in cerebellum, all individuals present a methylation level of approximately 75%, indicating biallelic methylation in some of the cells^6^. Results from Olsen et al. indicate that the bimodal methylation pattern is also lost from granulosa cells and that the methylation of the *nc886* locus in these specific cells might be associated with age^50^. Upon analysing somatic tissues not included in the previous study, we now report that the methylation pattern of breast, testis and prostate tissues are also distinct from what can be observed in most somatic tissues, including blood (Supplementary Figure 1Implications for future). Similarly to skeletal muscle, the bimodal methylation pattern cannot be observed in these tissues.

A small percentage of humans (1-6%) present intermediate methylation levels (20-40%) at the *nc886* locus and are thus chimeric of non-methylated and imprinted cells. We have shown that the intermediate methylation pattern in blood is not due to different cell type proportions^1^ and that it can also be observed across tissues^6^. Like the non-methylated and imprinted status, the intermediate status is also stable through time^1,13^. Furthermore, a few individuals (∼0.1% of the population) present methylation levels over 60%^1^ (Supplementary Figure 2), indicating that the paternal allele has also gained methylation in some proportion of the cells, thus expressing very low levels of nc886 RNAs. This implies that at least individual cells are viable without intrinsic expression of nc886 RNAs.

*nc886* is an evolutionally young gene. It can only be found in primates, guinea pigs and some members of the squirrel family^6,7^. In all the apes analysed previously (n=106), none of them presented a non-methylated epigenotype of *nc886*, indicating that the polymorphic imprinting of this locus is human-specific^27^. The centromeric CTCF site, which is also evolutionarily young, is relevant for the existence of the methylation pattern in the region, as only primates with an intact binding sequence of CTCF present the *nc886* imprint^27^.

### Methylation of *nc886* locus is associated with nc886 RNA levels

Regulation of gene expression through DNA methylation is a nuanced system. The best described mechanisms is the association between repressed gene expression and methylated CpG islands overlapping the transcription start site of a gene^51^. Methylation in the *nc886* DMR has been shown to regulate nc886 RNA expression in several *in vitro* settings, including 5-Aza-2′deoxycytidine treatment^11,28,52–54^. We and Treppendahl et al. have also shown that the intrinsic DNA methylation pattern in blood is associated with differences in nc886 expression levels^1,11^. In our data, the nc886-5p levels are increased two-fold in non-methylated individuals (both alleles permissive for expression) in comparison to imprinted individuals (only one allele permissive for expression). Further, individuals presenting intermediate DNA methylation levels in the *nc886* epiallele also present intermediate levels of the nc886 RNAs^1^ (Figure 1). Work by Park et al. describes how the methylation in the region leads to the formation of heterochromatin and the region being unavailable for RNA Pol III. They also suggest that, in open chromatin formation, MYC binds to an E-box upstream of the *nc886* gene and then interacts with RNA Pol III, enabling transcription^12^. In addition to this epigenetic regulation of nc886 expression, we have shown that genetic variation 100-200 kb downstream of the *nc886* gene is associated with the nc886 RNA levels^1^ (Figure 1B). As both the genetic profile and the DNA methylation pattern are stable in most somatic tissues and throughout an individual’s lifespan, there is already considerable physiological variation in nc866 RNA expression in healthy human population, which should be considered while investigating the levels of these RNAs in relation to morbidities.

### Establishment of the *nc886* imprint

Originally, Romanelli et al. suggested that the methylation pattern of *nc886* arises 4 to 6 days into embryonic development^42^. However, the results are based on cell lines derived from limited number of individuals. Reanalysing DNA methylation profiling data from the same cell lines (GSE52576)^55^ shows that the parthenogenetically activated oocytes and embryonic stem cells contain both non-methylated and imprinted cell lines (Supplementary Figure 3), indicating that these cell lines could actually be reflecting the intrinsic *nc886* methylation pattern of the oocyte. Immortalization and creation of iPSC have been shown to affect the methylation pattern of imprinted genes, including *nc886*, which should be taken into consideration when interpreting results from *in vitro* studies^6,12^. Our results from identical twins who have separated between days 1 and 3^6^ after fertilization and the analysis of oocyte data from Carpenter et al.^8^ also suggest that the methylation pattern is already established in the oocyte. Although DNA methylation profiling data is currently available from oocytes in many studies, the coverage of *nc886* DMR in bisulfite sequencing data is generally so low that no certain conclusion can be made. To some extent, this also includes the data of Okae et al.^56^ which the conclusions of Carpenter et al.^8^ and Jima et al. https://jb2.humanicr.org/^57^ are based on.

Although conclusive data is missing, current results are pointing in the direction of the *nc886* methylation pattern being established in the oocyte, in line with non-polymorphic maternally imprinted genes^58^. In mice, these imprints are established asynchronously during the growth and maturation of the oocyte in prophase II^59^. As the *nc886* polymorphic imprint is unique to humans, the establishment of the methylation pattern is difficult to study, as the growth phase of the oocyte lasts from early weeks after the birth of the woman to the formation of secondary follicle during menstruation, sometimes up to 50 years. The establishment of the intermediate and over-methylated patterns, on the other hand, likely happens during the de- and re-methylation of the embryonic genome, as identical twins separated before implantation can present methylation differences up to 17%^6^. This also fits the timing of the de- and re-methylation of the embryonic genome^60^ (Figure 2). Thus, we suggest that there are at least two different mechanisms contributing to the *nc886* methylation patterns observed in humans: the initial establishment of the pattern during oocyte maturation and then maintenance of this pattern during embryonic development.

**Figure 2.**
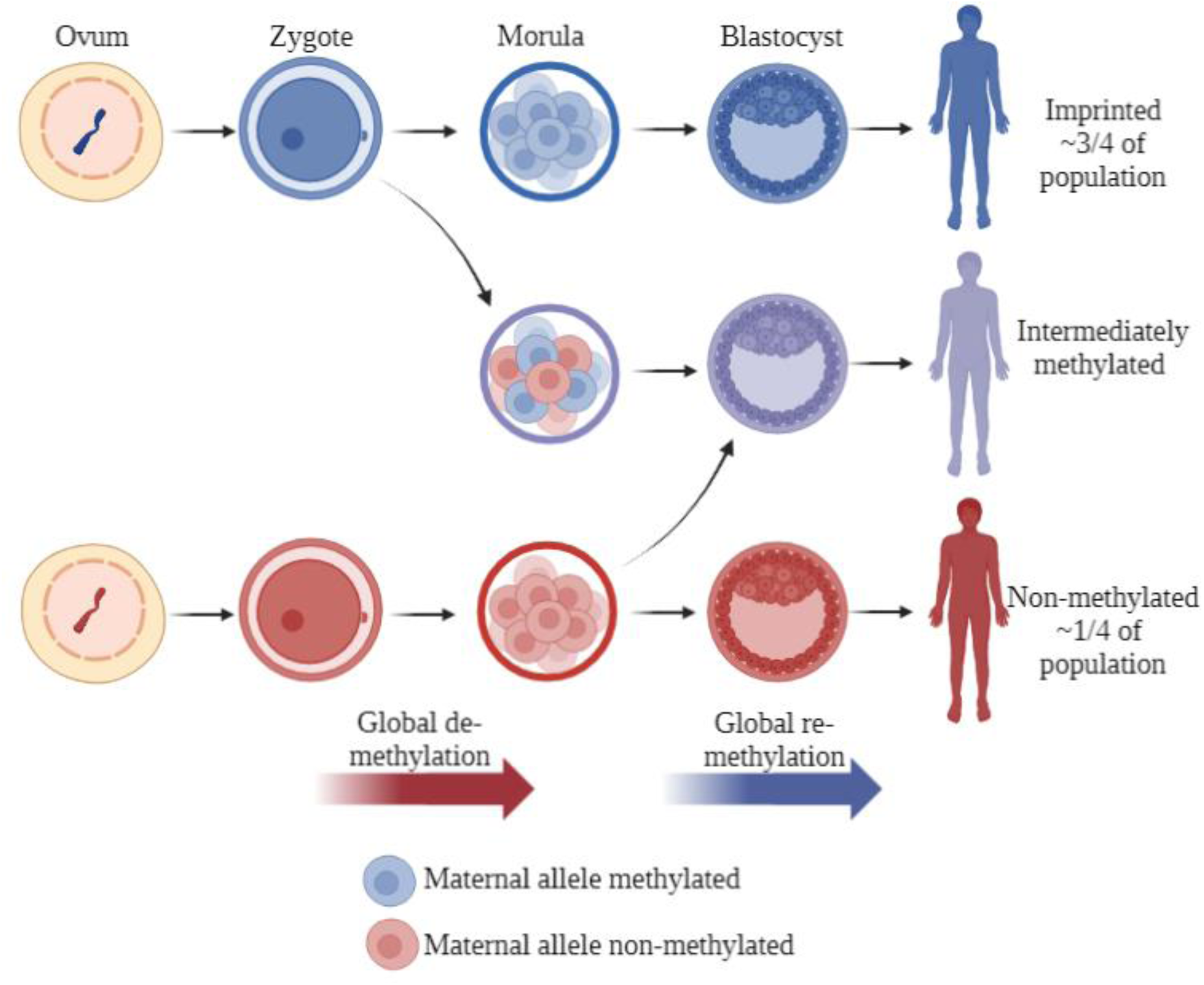
Establishment of the *nc886* methylation pattern. The methylation pattern of the non-methylated or imprinted *nc886* locus is suggested to be established during the maturation of the oocyte. We further hypothesize that the intermediate methylation pattern is caused by the sporadic loss of methylation during the global de-methylation of the embryonic genome or gain of methylation during the re-methylation. After implantation, the methylation pattern, and also rough portions of non-methylated and imprinted cells in intermediately methylated individuals, remain the same in the majority of somatic tissues. Made with BioRender.com.

One of the burning questions relating to *nc886* is what leads 25% of studied individuals to present the non-methylated epigenotype. Unlike with other polymorphically imprinted genes, targeted and genome-wide genetic analyses have failed to discover a genetic cause for the pattern^1,3,8,11,13^. Based on 30347 studied individuals from 32 datasets, we previously reported that in Caucasian singletons, the proportion of imprinted individuals varies less than 5%, ranging from 72.6 to 77.5%. When considering other ethnicities or populations consisting of twins, more variation could be observed^6^. In our previous study, populations consisting of individuals of African descent had a higher proportion of imprinted individuals (79.1% and 78.7%), whereas individuals of Asian descent had fewer imprinted individuals (68.2% and 65.8%), although the differences were not drastic, and the number of non-Caucasian populations was limited^6^. A smaller percentage of imprinted individuals (65.9 % n=82)) in an Asian population is also described in You et al.,^61^.

To provide more information on the subject, we investigated DNA methylation data from data repositories with a focus on non-Caucasian populations and were able to replicate our previous findings (Figure 3). When inspecting DNA methylation data from individuals originating from Africa, the highest percentage of imprinted individuals observed was 88% in ǂKhomani San, albeit that sample population is limited in size (GSE99029, n=57)^62^. The pattern is similar in populations of African origins from the US^63^, as well as other populations residing in Africa^64^. On the other hand, the percentage of imprinted individuals is lower in Asian populations, and the lowest percentage of individuals with imprinted *nc886* observed was 55% in populations in the Indonesian archipelago^65^ (Figure 3). It should be noted that within populations, differences can be seen between specific living location. We can’t thus rule out that the local genetic or environmental factors could affect the percentage of individuals with imprinted *nc886* locus, but also note that the observed differences could be explained by small numbers of individuals in the specific sub-populations (Supplementary Figure 4). As the establishment of the methylation pattern has not been associated with genetic variation ^1–3,5^, it is perplexing that the systematic variation in the percentage of the imprinted individuals was observed across populations. These results again highlight the need for more EWAS and mQTL analyses to be performed in non-Caucasian populations^66^.

**Figure 3.**
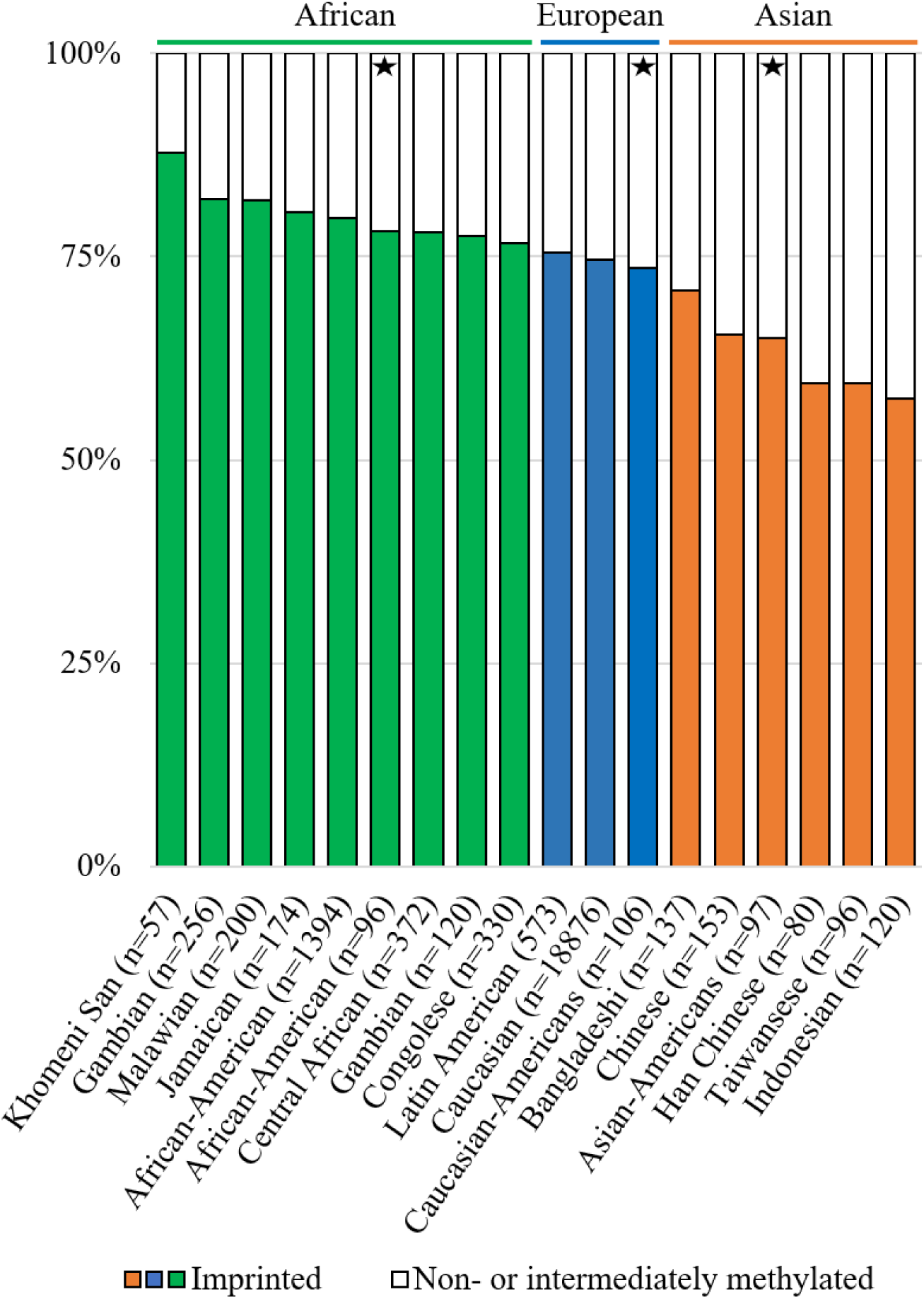
Percentages of imprinted (in colour) and non- and intermediately methylated individuals in *nc886* locus in population cohorts with ethnic origins in Africa (green), Europe (blue) and Asia (orange). In populations with ethnic origins in Africa, the percentage of imprinted individuals is higher and in those originating from Asia, the percentage is lower, than the 75% previously reported Caucasian populations^1^. Populations marked with a star are from the same sample series (GSE36369) and thus these populations are free of technical or sample collection bias when compared to each other. Data processing and thresholds for imprinted individuals are presented in Supplementary Materials and Methods and Supplementary Figure 2.

The prevalence of non-methylated individuals has been associated with maternal age^1,7,67^, the season of conception^7,13^, maternal nutrition^13^ or folic acid supplementation^68^, family socioeconomic status^1^ and maternal alcohol consumption^8^. Of these, only the association between maternal age and higher prevalence of non-methylated children has been shown in more than one cohort^7,13^. We further aimed to replicate the association between the season of conception and the prevalence of imprinted individuals. In the data set in which this was first reported, the analysis was performed with *nc886* methylation as a continuous variable^13^. Later, Carpenter et al.^7^ analysed the data as a categorical variable, but removed the intermediately methylated individuals as having “inconclusive” methylation pattern. We again re-analysed the data by dividing individuals into imprinted and others, to avoid losing individuals in a cohort with a limited size (n=120). While the finding linking the *nc886* methylation to the season of conception remains nominally significant, however, it does not replicate in a dataset with a similar study setting (GSE99863) (Figure 4A). All in all, more research with larger data sets is needed to understand if, and especially how, preconceptional or prenatal maternal traits associate with *nc886* methylation profile. It remains to be established whether these traits change the methylation status of the oocyte or embryo, or whether a non-methylated *nc886* region is beneficial for the success of fertilization or survival of the foetus in certain conditions.

**Figure 4.**
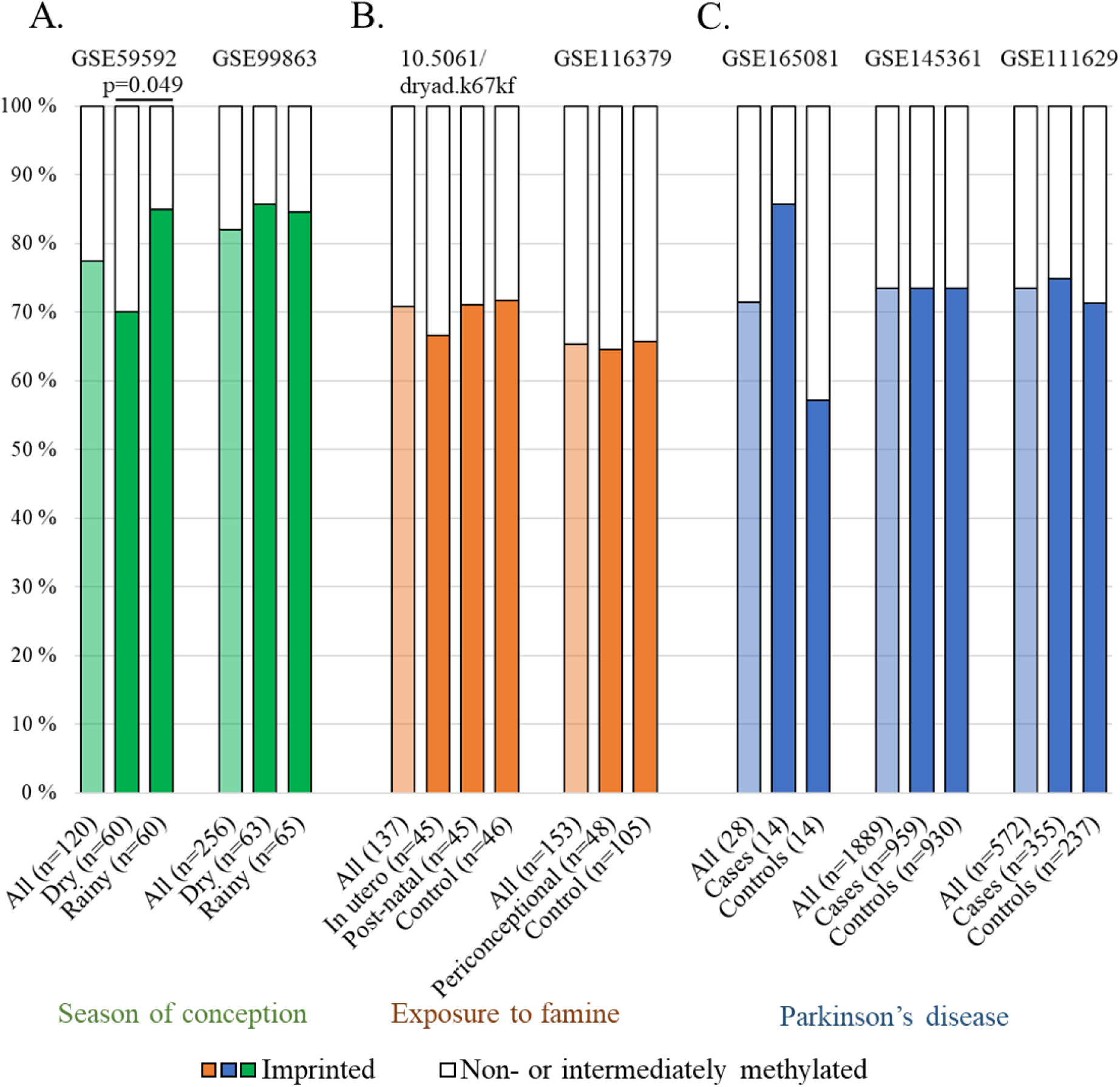
Re-analysing and attempting to replicate findings linking the *nc886* methylation pattern to A. the season of conception^13^ (GSE59592 and GSE99863), B. postnatal (10.5061/dryad.k67kf) and periconceptional (GSE116379) exposure to famine^14^ and C. Parkinson’s disease (GSE165081, GSE145361 and GSE111629). A. The original discovery by Silver et al. (GSE59592^13^) associating lower levels of *nc886* methylation to the season of conception remains statistically significant even after clustering individuals to imprinted and non-methylated or intermediately methylated (p=0.049), but this finding does not replicate in the other Gambian population cohort available. The significant association between post-natal exposure to famine (B) and Parkinson’s disease (C) disappear, when treating the *nc886* methylation pattern as a categorical variable, and are not replicated in data sets with similar or related study settings. Data processing and thresholds for imprinted individuals are presented in Supplementary Materials and Methods and Supplementary Figure 2.

If the proportions of *nc886* methylation status groups in a population are affected by mechanisms related to establishment and maintenance of the imprint, but also the chances of live birth in certain conditions, it would explain how *nc886* methylation pattern in populations could be affected by perinatal or periconceptional conditions while simultaneously being established already in the maturing oocyte. Further, the combined contribution of three distinct mechanisms (establishment, maintenance, and survival bias), could mask the potential genetic component in the pattern, thus explaining the differences between populations with different geological origins while simultaneously explaining the negative results observed in genetic analysis.

### Adulthood traits affecting *nc886* methylation status

The *nc886* methylation levels in blood have been shown to be stable for up to 25 years of follow-up in adulthood^1^ and from childhood to adolescence^13^. In contrast to these findings, a multitude of studies have reported, in cross-sectional settings, that methylation status in this locus is affected by outside exposures after birth or in adulthood, including post-natal famine^14^, adulthood exposure to pesticides or gases/fumes^69^, miner dust^70^, smoking^71^, wildfire-related fine particulate matter^72^, and long-term aircraft and railway noise^73^. One explanation for these conflicting findings is that the qualities of *nc886* methylation have not been taken into account and in these analyses, the methylation in the locus has been treated as a continuous variable, which can lead to false positive findings. We have now re-analysed the available dataset related to post-natal famine^14^ by categorising individuals based on the *nc886* methylation status. No difference was seen in the prevalence of imprinted individuals in those exposed to famine either pre- or postnatally in comparison to controls when re-analysing the data as categorical (Figure 4B). It should be noted that we were only able to analyse the whole data, not the selected subset utilised in the work of Finer et al. ^14^. Unfortunately, other datasets in which the above-mentioned associations were discovered are not publicly available, and thus could not be re-analysed with *nc886* methylation treated as a categorical variable. Furthermore, some of the findings are based on a very limited number of individuals and, for example, in the case of prenatal famine, similar studies have not been able to replicate the associations ^74–76^ (Figure 4). Thus, reanalysis of the data sets and replication are needed to rule out false positive findings.

### *nc886* and later life health traits

When treated as a categorical variable, the *nc886* methylation pattern has been associated with metabolic traits. We have shown that, in comparison to imprinted individuals, non-methylated individuals have higher insulin levels and glucose levels in childhood and adolescence, in childhood, boys have higher HLD and non-HDL cholesterol levels, and small children present higher adiposity ^1^. In line with this, van Dijk et al. showed that non-methylated children were in an increased risk of obesity, a trend which they replicated in a dataset consisting of 355 healthy young individuals (GSE73103)^77^. Maternal imprinted *nc886* locus has also been reported to be associated with increased risk of preterm birth (n=82)^61^, but this interesting discovery still needs to be replicated in a larger dataset.

Many publications report associations between the *nc886* methylation status and the risk of morbidities while treating the methylation in this region as a continuous variable. Lower methylation of *nc886* has been associated with increased risk of orofacial clefts^78^, IgA nephropathy^54^, and Parkinson’s disease^79^. When reanalysing the *nc886* methylation pattern as a categorical variable, it is clear that in the discovery cohort of IgA nephropathy (GSE72364, n=12^79^), four out of the six controls present non-methylated epigenotype and all of the six affected individuals are imprinted, generating the Δβ>0.3 (Supplementary Figure 2). With such a small population, the probability of having one group consisting only of imprinted individuals is 18%, and the nominally significant results would be obtained with having 3 out of 6 individuals non-methylated in the other group. Similarly, in data of Henderson et al. (GSE165083^79^), the noticeable, but statistically non-significant difference in the *nc886* methylation status groups between Parkinson’s disease cases and controls (imprinted vs others in chi-square test p=0.09, n=28) is generated by 8 vs. 12 imprinted individuals in each group (Supplementary Figure 2). Furthermore, no difference can be observed in the *nc886* methylation patterns in Parkinson’s patients and controls in GSE145361^80^, with 1889 individuals or GSE111629, with 572 individuals^6,81^ (Figure 4E).

Similar issues can be seen when reporting the associations of nc886 RNA expression and phenotypes in small sample settings. For example, Miñones-Moyano et al. report elevated nc886-5p levels in the brains of Parkinson’s disease patients in comparison to controls (n<40 per setting). A clustered expression pattern can be observed in the amygdala, the frontal cortex and the substantia nigra of Parkinson’s patients with patients at motor stages of the disease (in Miñones-Moyano et al. ^16^ Figure 1). As the nc886 RNA levels are strongly regulated by the methylation pattern, their results could be caused by the uneven distribution of imprinted and non-methylated individuals in cases and controls. In support of this, the aforesaid connection to Parkinson’s disease cannot be detected in the cerebellum, where the bimodal methylation pattern cannot be detected^6,16^. Similarly, it would be interesting to see whether the associations between nc886 RNA levels and IgA nephropathy would remain significant after taking into account the stable regulators of the RNA levels^82^. A reanalysis with the methylation and genetic regulators included in the model would be warranted to confirm these results.

### *nc886* and cancer

Changes in *nc886* methylation level and RNA levels are widely reported in cancer. nc886 RNAs have been found to be upregulated in cervical cancer^83–85^, breast cancer^84,86^, high grade bladder cancer^87^, high grade prostate cancer^88,89^, multiple myeloma^90^, endometrial cancer^31^and renal carcinoma^91^, while down regulation has been reported in cholangiocarcinoma^29^, oesophageal cancer^52,92^, prostate cancer^93,94^, ovarian cancer^95^, breast cancer^96^, thyroid cancer^97,98^, small-cell lung cancer^99^, squamous cell lung carcinoma^100^ and oral squamous cell carcinoma^101^. Notably, in breast and prostate cancers, both directions are reported. Several molecular mechanisms could explain the changes in *nc886* transcription in cancer. The expression of RNA pol III transcribed genes, such as *nc886* has been reported to be up regulated in cancer. The transcription of *nc886* is also activated by transcription factor MYC, the levels of which are often also up regulated in cancerous cells^12^, theoretically leading to even higher rates of expression of nc886 RNAs. On the other hand, hypermethylation of the *nc886* locus has been reported in several types of cancers^42,53^, leading to the formation of heterochromatin and repression of the transcription^12^.

Knocking out or down the nc886 RNA expression has been shown to both induce apoptosis or suppress proliferation^25,31,83–86,89,91,102^ and promote cell division^103^. Similarly, overexpression of these RNAs has been shown to have both growth promoting^83,84,89,91,104^ and restricting^9,52,93,97,101,105–107^ properties. Elevated levels of the nc886 RNAs or lower levels of the DNA methylation in the region have also been linked to both worse ^28,89,99,108–110^ and better^11,52,94,103,105,106,111,112^ prognosis of the malignant disease. The issues of investigating the roles of alleged miRNAs in cell cultures with over-expression or knocking out has been described in detail by Lee et al., emphasising the selection of suitable methods to produce a natural nc886 molecule in physiological concentrations and correct methods in evaluating the success of knock-down^10^. In the case of *nc886*, the natural variation in the stable regulators of the RNA expression should also be taken into account. The stable methylation pattern and the genetic variation in the proposed enhancer area of the gene can lead to substantial disease status-independent differences in RNA levels^1^, which can lead to false positive findings when studies are performed in small sample sets or selected cell lines.

Changes in the methylation pattern of the *nc886* locus have also been linked to cancer^42,53,113^. However, the majority of these studies do not take into account the bimodal distribution of normal tissue or that the changes in the DNA methylation pattern lead into different consequences in relation to the original methylation status. If the methylation status increases by 20%, it will lead to very different results depending on whether the inherent methylation level was 0% or 50%. In those studies, where the original methylation pattern has been considered, both hyper- and hypomethylation have been reported^42,53,113^, with hypermethylation being associated with poorer survival and increased risk of tumour recurrence^113^. Also, the innate DNA methylation status of *nc886* locus has been shown to predict the future risk of prostate^4,5^, breast^5,41^ and pancreatic cancer^114^, although the study by Wang et al. treats *nc886* methylation as a continuous variable and reports rather inconsistent results^114^. These studies suggest that individuals with non-methylated *nc886* might be at increased risk of developing cancer, which would, at least in theory, be in line with the suggested growth promoting role of maternally imprinted genes, such as *nc886*, during foetal development^115^.

Ignoring the special qualities of *nc886* methylation can also hide biologically relevant results. As an example, when comparing the methylation levels in the region from clear cell renal cell carcinoma (GSE61441) and adjacent normal tissue, a statistically non-significant (p=0.06) hypomethylation can be detected in the tumours. When comparing the within individual differences between normal tissue and tumour, 65% (30/46) of the tumours show more than 2% hypomethylation while 13% (6/46) present hypermethylation over 2%, suggesting a pattern of hypomethylation, but also stark differences between individual tumours. Intriguingly, hypomethylation can also be seen individuals presenting low methylation levels in healthy tissue (Figure 5).

**Figure 5.**
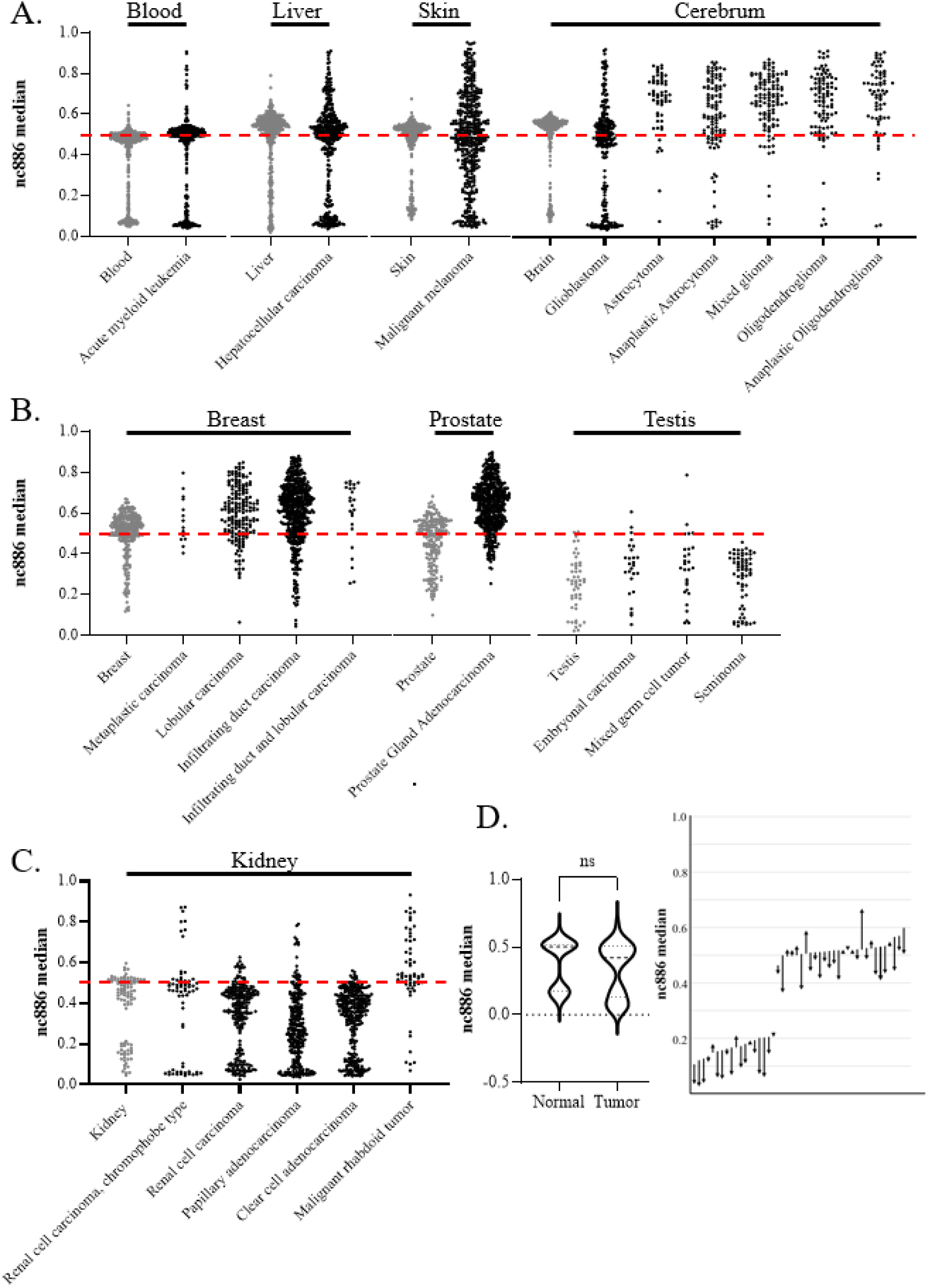
(A-C) The median methylation level of *nc886* in tumour samples (in black) from The Cancer Genome Atlas Program (TCGA) and from corresponding healthy tissues (in grey). A) In some cancer types, the original bimodal methylation pattern is faithfully or near-faithfully maintained (acute myeloid leukaemia and hepatocellular carcinoma) while in others the pattern is completely lost (malignant melanoma). In tumours from the cerebrum (excluding glioblastoma), a systematic hypermethylation can be observed. B) The bimodal methylation pattern of *nc886* is missing already from the healthy breast, prostate and testis samples, with breast and prostate cancer demonstrating hypermethylation, while the pattern in the tumours from the testis corresponds to the hypomethylated unimodal pattern observed also in the healthy tissue. C) Interestingly, renal carcinomas are observed to present hypomethylation in comparison to the healthy tissue, while some malignant rhabdoid tumours of the kidney are hypermethylated. D) When analysing *nc886* median methylation from the healthy adjacent tissue and tumour from clear cell adenocarcinoma (GSE61441)^133^, a non-significant minor hypomethylation in the tumours can be observed, but while analysing the changes in individual level more systematic hypomethylation pattern can be detected. The start of the arrow presents the methylation level of the healthy tissue, and the end of the arrow the methylation level of the tumour, with each arrow presenting one individual. Data processing and thresholds for imprinted individuals are presented in Supplementary Materials and Methods and Supplementary Figure 5.

We retrieved data from The Cancer Genome Atlas Program (TCGA) to investigate the *nc886* methylation pattern in cancerous tissue and discovered that the changes in methylation are cancer type specific. While in some cancers, such as acute myeloid leukaemia, the methylation pattern remains mostly unaltered, in some, like malignant melanoma, the bimodal methylation pattern observed in healthy skin is completely lost (Figure 5). In the majority of the cancer types, some degree of hypermethylation and loss of control of the methylation pattern can be observed (Figure 5). More systematic hypermethylation, in comparison to the bimodal pattern in blood, can be observed in brain cancers, excluding gliomas, prostate and breast cancers. Hypomethylation in comparison to healthy tissue can be observed in renal carcinomas. Both healthy and cancerous testes present lower methylation levels than blood, and an absence of the bimodal methylation pattern. Similarly, in both healthy breast and prostate tissue the bimodal methylation pattern is not present, as majority of individuals present methylation levels around 50% and in tumours from these tissues, hypermethylation can be seen (Figure 5)

It will be interesting to study, whether it is the methylation level itself, degree of hypo- or hypermethylation, or the loss of the established pattern that is associated with cancerous development and future prognosis. One can hypothesise that the loss of imprint in this region could be a sign of epigenetic instability, thus associated with a poorer prognosis, regardless of the direction of change in the methylation levels. However, the original methylation pattern of the individual and tissues of origin, as well as the proportions of the methylation status groups in a population, should always be taken into account when interpreting the obtained results. For example, the findings from a cross-sectional study suggesting that human papilloma-virus is associated with similar changes in *nc886* methylation profile in both the cancerous and the healthy tissue could also be explained by different proportions of imprinted individuals in case and control groups at baseline ^102^. Due to the peculiar nature of *nc886* methylation, longitudinal analyses would be exceptionally beneficial to determine its role in cancer progression.

## Conclusions

*nc886* has been shown to be associated with both periconceptional conditions and adulthood health traits, but many of these results do not replicate in independent cohorts. Results obtained from small cohorts especially warrant reanalysing and replication. Current knowledge indicates that the DNA methylation pattern in the *nc866* locus is established during the oocyte maturation^6,8^, is stable during life and in the majority of somatic tissues, but is associated at least with maternal age at birth^1,7^. In this case, it should be contemplated whether *nc886* can be considered to mediate the DoHAD hypothesis, as the methylation status as such is not affected by environmental conditions. However, if the methylation pattern in a population is shaped by pregnancy success, *nc886* could mediate genetics-independent adaptation of the population to the surrounding conditions.

Approximately 75% of Caucasians present an imprinted *nc886* locus, while populations with African origins present a higher percentage of imprinted individuals, and those originating from Asia have systematically lower numbers of imprinted individuals. Genetic and epigenetic analyses with more diverse backgrounds are needed to understand this pattern. In previous works, *nc886* methylation has been associated with metabolic traits and immortalization and carcinogenesis have been shown to alter the inherent DNA methylation pattern^12^. We demonstrate here that the changes in the methylation pattern are cancer-type and tissue-of-origin specific. It is, however, still unclear, what the role of nc886 RNAs in the development of malignancy is. Similarly, the true nature of nc886 RNAs is still under debate. As the hairpin structure suggested to bind PKR is formed by nucleotides 46-59, and the short RNAs are suggested to be processed from nucleotides 1-24 and 80-101, both hypotheses could potentially be correct.

Great inherent variation caused by the stable genetic and epigenetic regulators in the nc886 RNA expression within populations can lead to wrong interpretations, when assuming that all differences between cases and controls are due to the studied condition. Furthermore, the binomial methylation pattern of *nc886* warrants post-hoc analyses every time the region is discovered in an EWAS analysis. These notions can be generalized to study of DNA methylation and expression of gene products regulated via DNA methylation. Genetic variation and other features, such as sex, can contribute to methylation levels that are categorical, rather than continuous. Especially in studies with small datasets, it would be important to inspect the distribution of identified methylation sites. Further, even in continuous features, there is potentially physiological variation in levels of the measured epigenetic profile, not caused by condition in question. Even though *nc886* codes for only a few peculiar RNAs and is located in an atypical locus, the contradicting results presented here highlight that in this era of genome-wide bioinformatic analyses and vast amounts of data, researchers should take time to study their top findings further, to avoid reducing science to mere reporting of statistically significant values.

## Implications for future studies

1. The DNA methylation pattern in the *nc886* locus should be treated as a binomial variable to avoid reporting false positive findings due to the random distribution of non-methylated and imprinted individuals in cases and controls.
2. While studying the DNA methylation pattern in cancer, hypo- and hypermethylation of *nc886* should be reported only when the intrinsic DNA methylation pattern has been taken into account in the analysis.
3. When investigating the associations between the nc886 RNA levels and phenotypes, the stable genetic and epigenetic regulation pattern of the individuals or cell lines should be taken into account, to avoid false positive results from the uneven distribution of the stable regulatory profiles of nc886 RNAs in small case-control settings.

## Open questions relating to *nc886*

1. How and when is the *nc886* DMR established and do the preconceptional/prenatal conditions modulate the DNA methylation pattern? Does the non-methylated methylation pattern, for example, provide a survival advantage to the foetus in non-optimal pregnancy conditions? What causes the distinct patterns of the *nc886* methylation status in cohorts from different geographical origins? How are the intermediate and over-methylated *nc886* methylation patterns established?
2. Is the functional form of nc886 RNAs the 108nt long hairpin structure binding to proteins, the two short RNAs produced by Dicer, acting in a miRNA-like manner, or both? What is the function of nc886 RNA(s) in physiological conditions?
3. Does *nc886* have a causal role in carcinogenesis, or are the changes in DNA methylation pattern and RNA expression consequences of epigenetic instability?

## Data availability statement

All new analysis performed have utilized data freely available or available upon reasonable request. For different geographic origins data from Khomeni San (GSE99029^62^), Gambian (GSE99863^116^), Malawian and Jamaican (GSE112893^117^), African American (GSE210255^118^), Multi-ethnic American (GSE36369^63^), Central African (https://ega-archive.org/; EGAD00010000692^64^), Gambian (GSE59592^13^), Congolese (GSE224363^119^), Latin American (GSE77716^120^), Caucasian (summary statistics^6^), Bangladeshi (http://datadryad.org/, doi:10.5061/dryad.k67kf14), Chinese (GSE116379^76^) Han Chinese (GSE201287^121^), Taiwanese (GSE78904^122^) and Indonesian (https://figshare.com, 10.26188/5e00abd72f581^65^) cohorts were utilized. Re-analysing and attempting to replicate previous associations of *nc886* methylation pattern and phenotypes, the follow data seta were used; parthenogenetically activated oocytes and embryonic stem cells (GSE52576^55^); for season of conception GSE59592 and GSE99863; for postnatal and prenatal exposure to famine (http://datadryad.org/, doi:10.5061/dryad.k67kf) and GSE116379 and for the presence of Parkinson’s disease (GSE165081^79^, GSE145361^81^ and GSE111629^123^). Methylation data from tumours was downloaded from The Cancer Genome Atlas Program (TCGA) and the reference healthy tissues data was utilized as follows: brain (GSE72778^124^), blood (GSE40279^125^), liver (GSE61258^126^ and GSE180474^127^), skin (GSE90124^128^), prostate (GSE76938^129^ and GSE213478^130^), breast (GSE88883^131^, GSE101961^132^ and GSE213478^130^), testis (GSE213478^130^) and kidney (GSE61441^133^ and GSE213478^130^).

## Acknowledgements

We wish to thank Daria Kostiniuk (MSc) and Netta Pihlanen for their contribution for the manuscript. We would also like to thank all the researchers, who have published on *nc886* and especially those who have produced the data utilized here and made it freely available or provided it upon request.

## Author contributions

ER conceptualised and wrote the first draft of the manuscript, provided data and figures, and reviewed and edited the manuscript. SR provided data and revised the manuscript. SM conceptualised the manuscript and reviewed and edited it.

