## Supplementary File for "Non-coding 886 (*nc886*/*vtRNA2-1*), the epigenetic odd duck – implications for future studies"

- Supplementary Figure 4. ........ 4
- Supplementary Figure 5. ........ 8

### Supplementary methods

Majority of the data sets utilized in this paper have been accessed through the Gene Expression Omnibus (GEO https://www.ncbi.nlm.nih.gov/geo/) or The Cancer Genome Atlas Program (TCGA, https://www.cancer.gov/ccg/research/genome-sequencing/tcga) or have been otherwise freely available. Permission to utilize data was requested from the contact person for the Central African population (https://ega-archive.org/; EGAD00010000692). All data sets were downloaded as processed data. If data was available as M values, these were transformed to beta values (Beta = 2^M/(2^M + 1)).

11 CpG sites (cg07158503, cg11608150, cg06478886, cg04481923, cg18678645, cg06536614, cg25340688, cg26896946, cg00124993, cg08745965, cg18797653) were extracted and median values of these were calculated. All data sets were plotted (Supplementary Figures 1-2) and suitable threshold to detect imprinted individuals was selected for population cohorts. In majority of the cohorts, threshold of 0.4 was utilised to subset the population. With GSE99863, GSE36369, GSE224363, GSE165081 and GSE145361 the threshold was set to 0.35, as visual inspection revealed that the majority of the imprinted individuals had methylation levels between 0.4 and 0.5.

Percentages of individuals presenting imprinted *nc886* status were calculated for whole populations and in some cohorts also in subpopulations or in cases and controls separately. When comparing the prevalence of imprinted individuals in cases and controls chi-squared test was utilized.

For data from TCGA and healthy tissues corresponding to the tissues of origins of the cancers, no thresholds were set, but the data was visually investigated. For the control tissue for the cancers of cerebrum (GSE72778) cerebellar samples were removed, as the data presented as clear outliers (Supplementary Figure 1).

### Supplementary figures

**Supplementary Figure 1.** Scatter plots of the median methylation level of the 11 probes of *nc886* locus from the heathy tissues utilized as controls for the Figure 5. Cerebellar samples were removed from GSE72778, as they behave differently than regions of cerebrum, as previously reported[1].

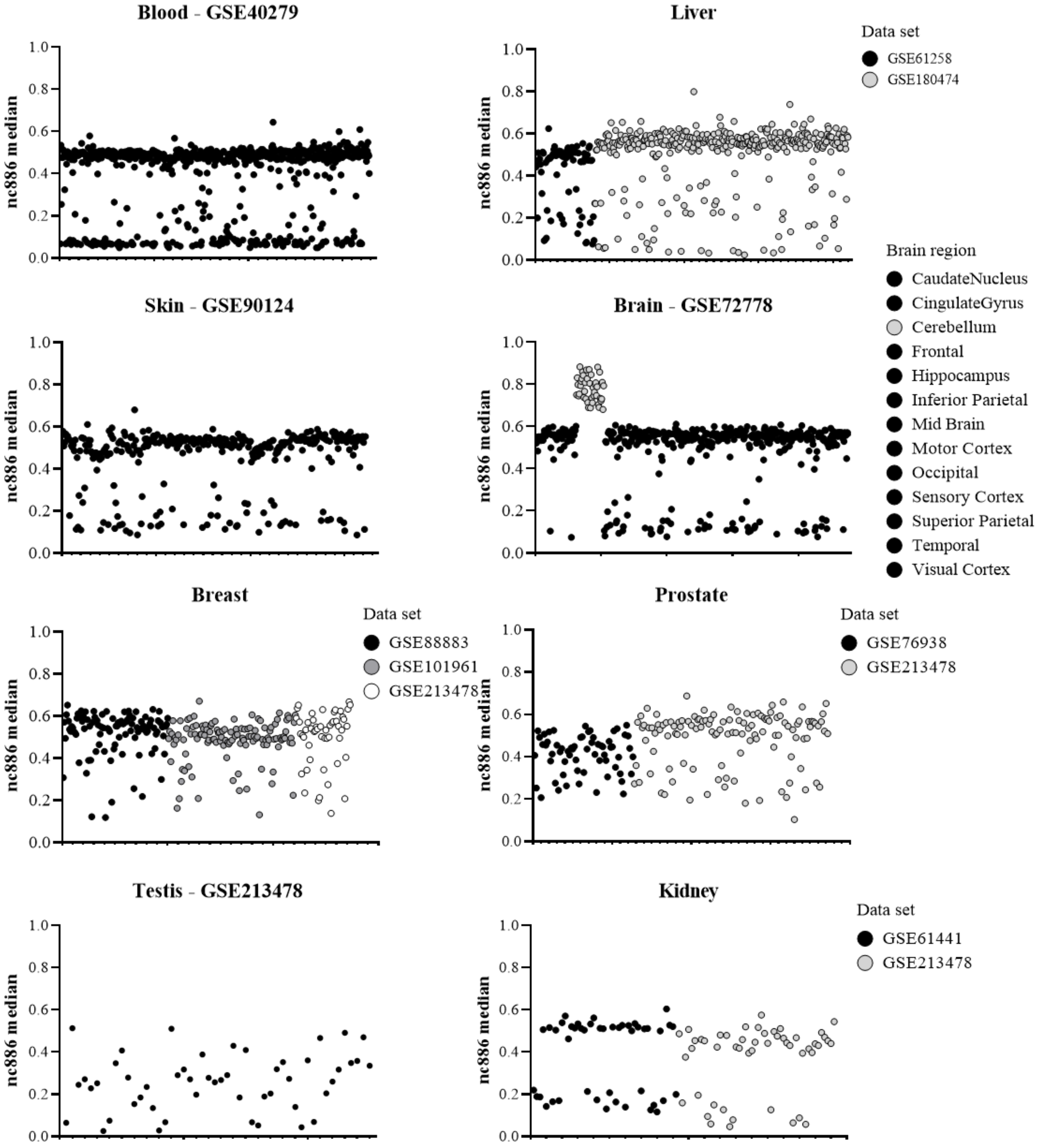

**Supplementary Figure 2.** Scatter plots of the median methylation level of the 11 probes of *nc886* locus from all the population cohort utilized in Figure 3. Data set specific *nc886* median beta-value thresholds for imprinted individuals have been illustrated with a dashed line.

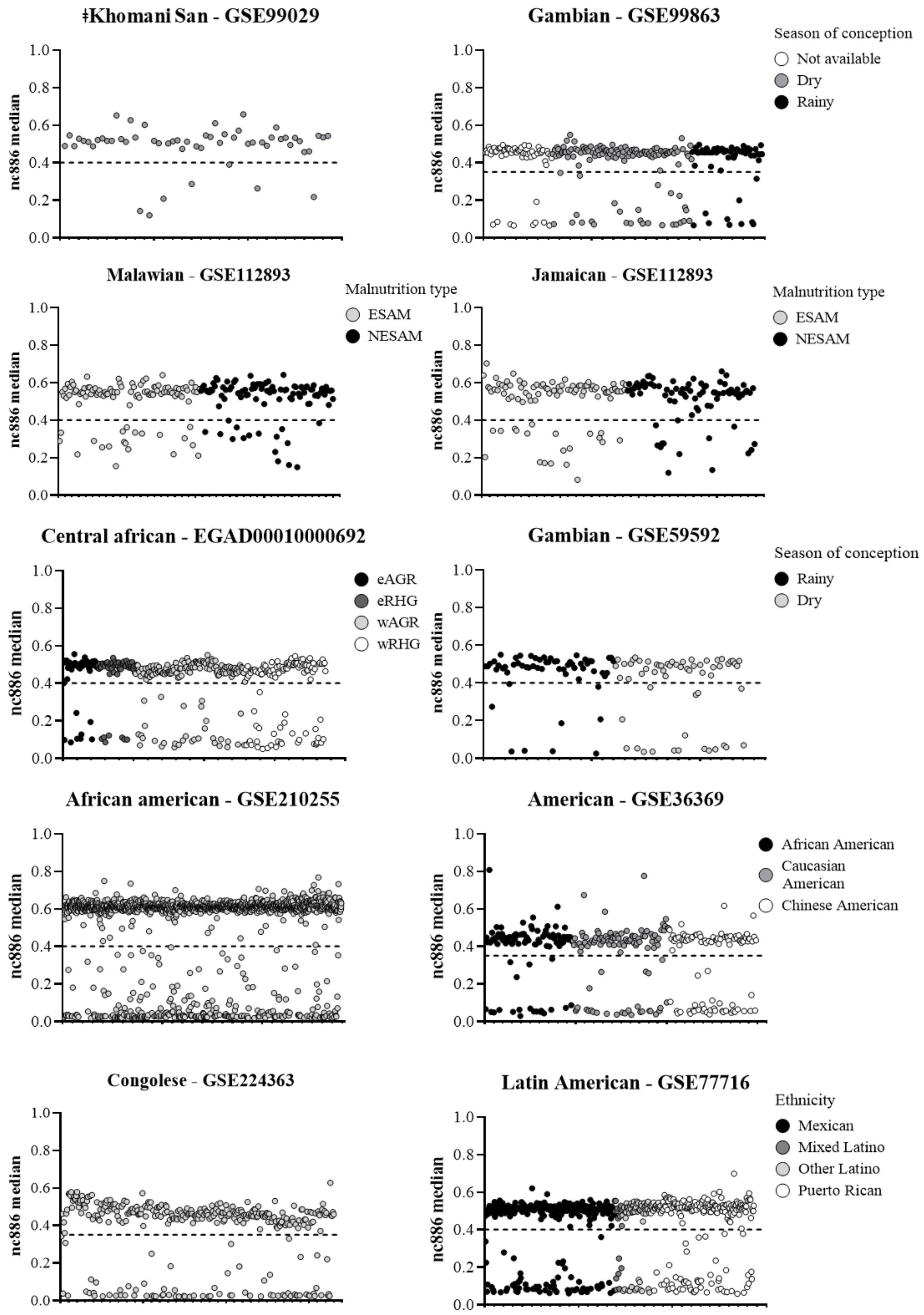

**Supplementary Figure 2. continues.**

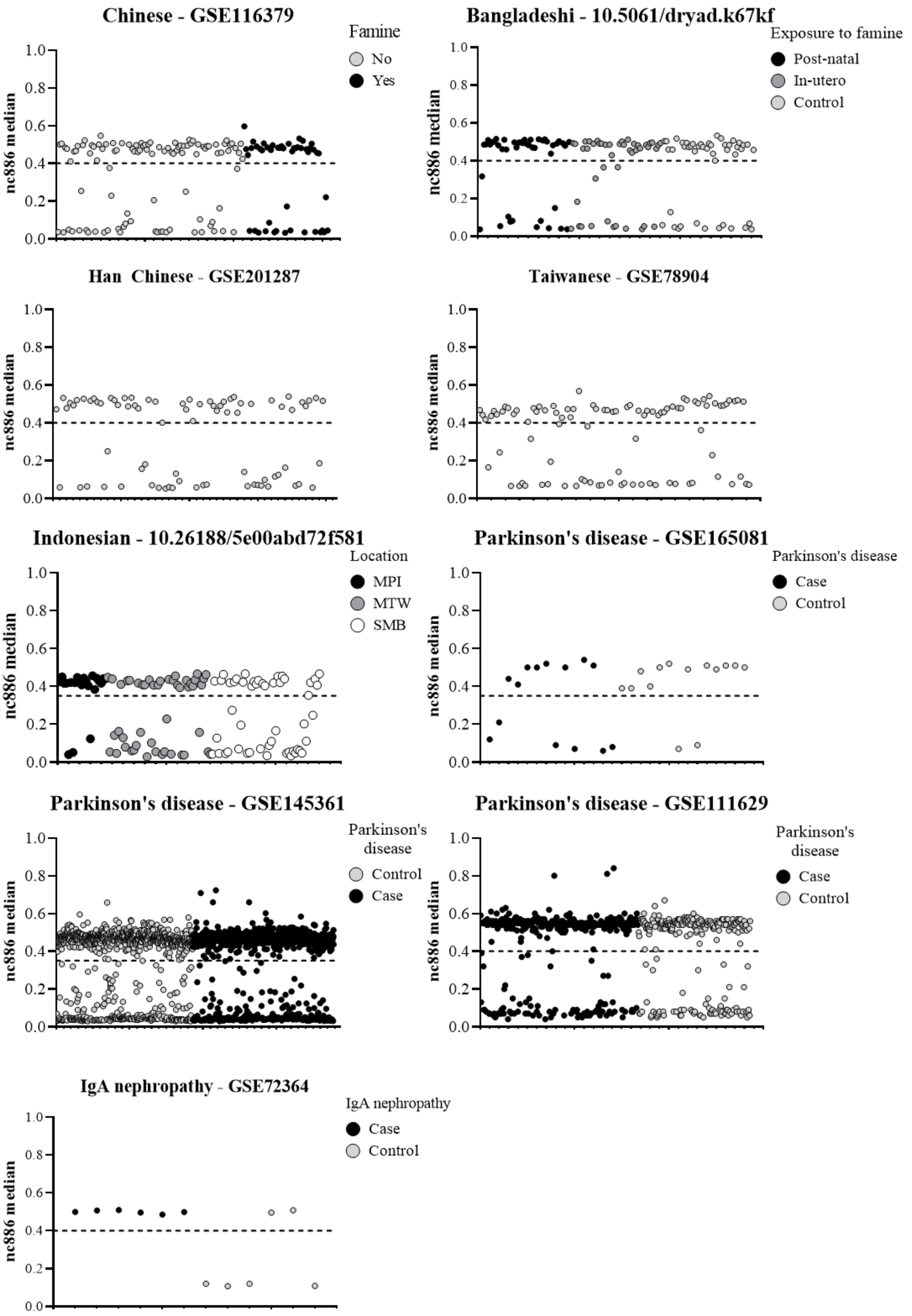

**Supplementary Figure 3.** Median methylation levels of *nc886* locus from GSE52576[2].Presented are four cell lines created from parthenogenetic activated oocytes (in grey) and from four human embryonic stem cell lines (in black). Both groups precent individuals cell lines with imprinted *nc886* locus (methylation ~50%) and those with very low methylation pattern (methylation <10%).

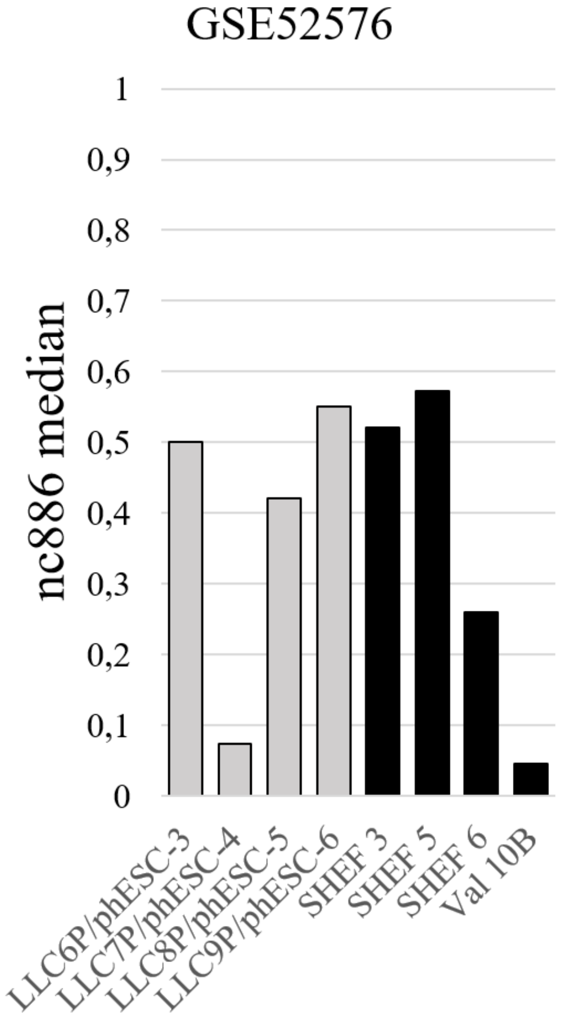

**Supplementary Figure 4.** Percentage of imprinted individuals from the subpopulations of the A. Central-African (https://ega-archive.org/; EGAD00010000692[3]) and B. Indonesian (https://figshare.com, 10.26188/5e00abd72f581[4]) cohorts. In the Central-African cohort, the western rainforest hunter-gatherers (wRHG) and in the Indonedian cohort the Korowai (MPI) stand out from the other sub-populations. The Indonesian population includes only 20 participants from Korowai and the distinct pattern can be due to chance in a limited population. However, the Central-African population includes 114 western rainforest hurter-gatherers, implying that local genetic or environmental factors cannot be ruled out as causal factors in the percentage of imprinted individuals.

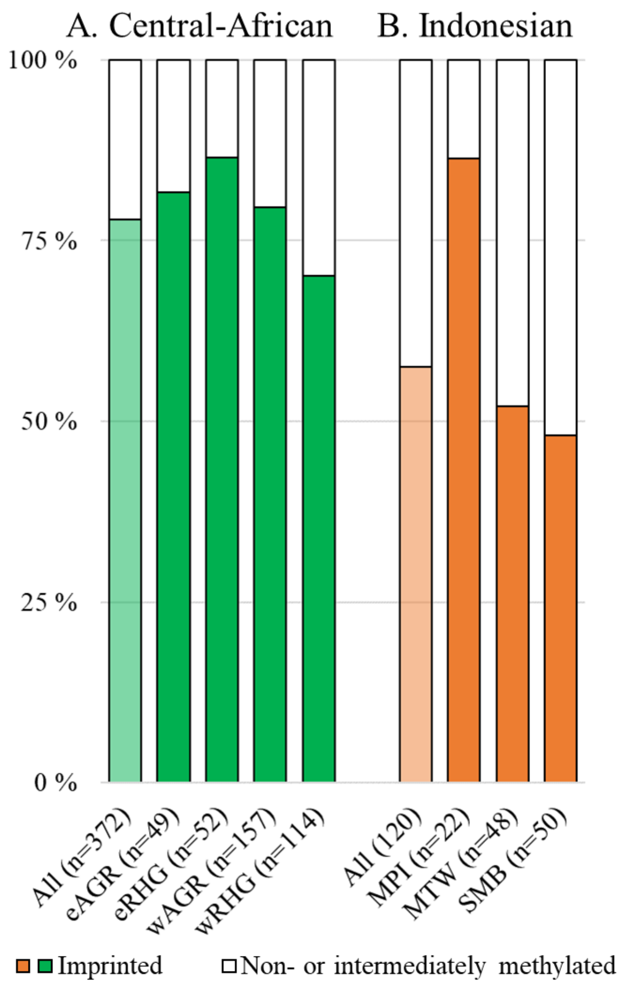

Abbreviations: eAGR=Eastern agrarian lifestyle, eRHG=Eastern rainforest hunter gatherer, wAGR=Western agrarian lifestyle, wAGR=Western rainforest hunter gatherer, MPI=Korowai, MTW=Mentawai, SMB=Sumba

**Supplementary figure 5.** Scatter plots of the median methylation level of the 11 probes of *nc886* locus from tumour samples utilized in figure 5. The data has been obtained from The Cancer Genome Atlas Program (TCGA) (https://www.cancer.gov/ccg/research/genome-sequencing/tcga).

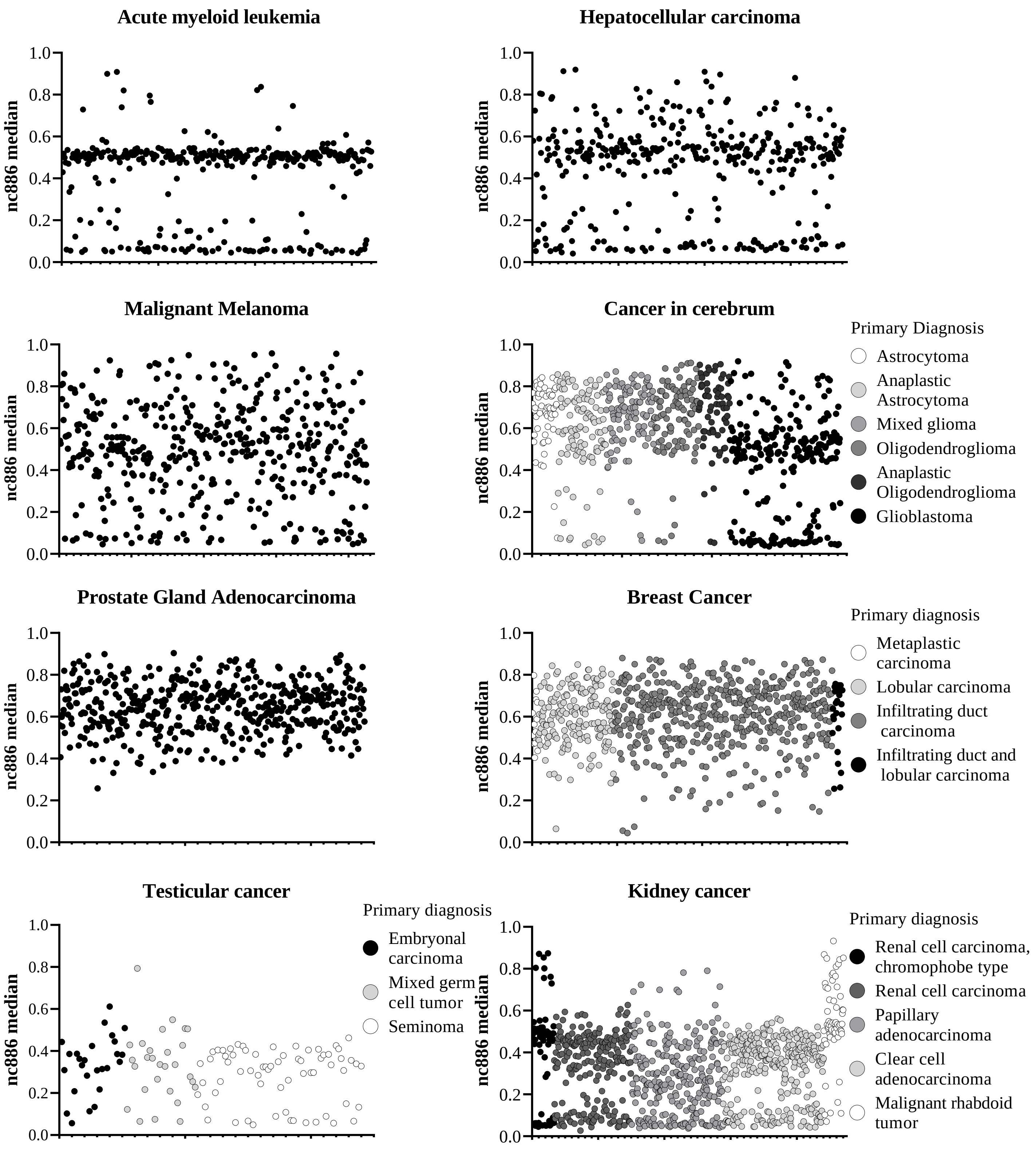
